# Prediction Interval Ranking Score: Identification of Invariant Expression from Time Series

**DOI:** 10.1101/482794

**Authors:** Alexander M Crowell, Jennifer J. Loros, Jay C Dunlap

## Abstract

**Motivation:** Identification of constitutive reference genes is critical for analysis of gene expression. Large numbers of high throughput time series expression data are available, but current methods for identifying invariant expression are not tailored for time series. Identification of reference genes from these data sets can benefit from methods which incorporate the additional information they provide.

**Results:** Here we show that we can improve identification of invariant expression from time series by modelling the time component of the data. We implement the Prediction Interval Ranking Score (PIRS) software, which screens high throughput time series data and provides a ranked list of reference candidates. We expect that PIRS will improve the quality of gene expression analysis by allowing researchers to identify the best reference genes for their system from publicly available time series.

**Availability:** PIRS can be downloaded and installed with dependencies using ‘pip install pirs’ and Python code and documentation is available for download at https://github.com/aleccrowell/PIRS.

## Introduction

The identification of invariantly expressed reference genes underlies many studies in molecular biology. With the advent of widespread qRT-PCR studies, several tools were developed to identify constitutive reference genes from these experiments Vandesompele *et al*., 2002; Andersen *et al*., 2004; Pfaffl *et al*., 2004. More recently, several tools have been developed for identifying stably expressed reference genes from high throughput experiments Hruz *et al*., 2011; Carmona *et al*., 2017. Unfortunately, some reference genes thought to exhibit constitutive expression have later been found to be dynamically expressed in time series. GAPDH in particular was employed widely as a reference gene - and is still used in some cases - in spite of its circadian expression in many model organisms including mice Shinohara *et al*., 1998; Kosir *et al*., 2010; Pizarro *et al*., 2013. High quality time series expression data sets are available for a wide array of model organisms, yet reference genes are often still identified by analyzing small numbers of genes with tools not designed for time series Mughal *et al*., 2018; Jain *et al*., 2018. We seek to address this issue by leveraging the structure of time series experiments in order to better identify invariant expression.

PIRS first screens out clearly dynamic expression profiles using an ANOVA. The algorithm then looks for invariant expression both within and between time points for an expression profile by comparing mean expression to the prediction intervals of a linear regression. We report the results of simulations comparing the performance of PIRS to the more commonly used selection using the standard deviation/coefficient of variation. In these simulations, PIRS out performs methods which do not model the time component of the data. PIRS has previously been used to identify reference genes in *N. crassa* Hurley *et al*., 2015.

## Methods

### Algorithm Design

PIRS first performs an ANOVA on each expression profile to determine if any time points are differentially expressed and screens them out. This screening step both improves precision by removing clearly unwanted profiles from analysis and improves running time by reducing the number of profiles for which a score must be calculated. The algorithm then performs a linear regression of expression profiles *y* against the *n* time points *x* and calculates prediction intervals for these regressions. With prediction intervals:

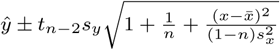

where:

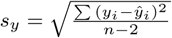

and *t*_*n*-2_ is the 1 − *α*/2 ppf of the student’s t-distribution with *n –* 2 degrees of freedom.

The absolute difference between the mean expression of the profile and the prediction intervals is then calculated for all time points. The final ranking score is derived by summing these differences and dividing by the mean expression. A truly constitutive expression profile would be expected to remain close to its overall mean across all observations and also to have relatively low variance around that mean. We would therefore expect the prediction intervals in this case to remain close to the overall mean for all observations. In contrast, we would expect a linear expression profile to have narrow prediction intervals which are far from the mean at most time points and a circadian expression profile to have prediction intervals centered on the mean but which are nonetheless far from it because they are broadly spaced (Fig S1 a).

### Simulation Studies

In order to assess the effectiveness of PIRS at distinguishing constitutive from dynamic expression profiles, we generated simulated data sets and benchmarked PIRS against a standard deviation based method. We compared performance on both circadian and linear trends with gaussian noise added to all profiles. In both cases, 20% of profiles were truly invariant. In both tests, PIRS markedly outperformed the standard deviation based method (Fig S1 b and c).

## Conclusion

When analysing time series data, the replicate and time point structure provide valuable information. PIRS provides improved results when screening time series data for invariant expression by incorporating this information and tailoring the analysis to the data. With PIRS, researchers can effectively identify reference genes from high throughput time series data sets in their model system. By identifying better reference genes, PIRS enables better down stream experiments.

## Supporting information

## Funding

Funding for AMC was provided by NIH grants R01GM083336 and 1R35GM118022 to JJL, 1R35GM118021 and 1U01EB022546 to JCD, and through an Albert J. Ryan Fellowship.

