## Supplementary material for "Prediction Interval Ranking Score: Identification of Invariant Expression from Time Series"

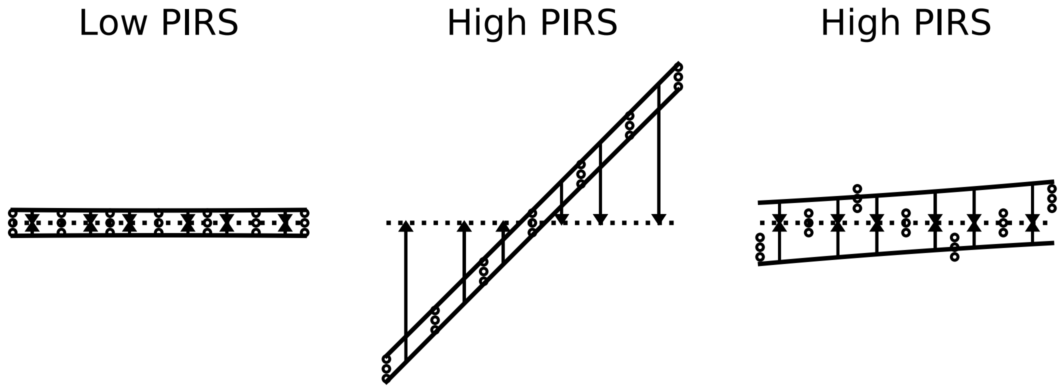


(a) Example PIRS Calculations

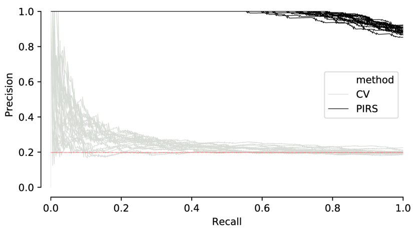


(b) Precision Recall for Linear Trends

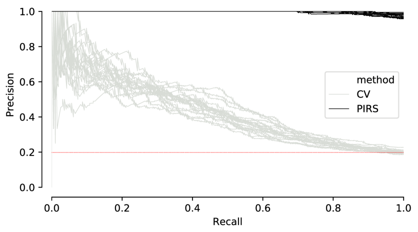


(c) Precision Recall for Circadian Trends

Figure 1: PIRS: Description and Performance
(a) Example PIRS calculations. Circles indicate data points, dashed horizontal lines indicate mean of all data, solid lines show prediction intervals and arrows show the difference between prediction interval and mean. (b) PIRS performance on linear trends. Dark lines show precision/recall for PIRS, light lines show precision/recall for the standard Coefficient of variation method. Dashed horizontal line represents a random classifier. (c) Results for Circadian trends. Dark lines show precision/recall for PIRS, light lines show precision/recall for the standard Coefficient of variation method. Dashed horizontal line represents a random classifier
